# SIRT1 limits neuronal fate following DNA damage

**DOI:** 10.1101/2025.05.19.654885

**Authors:** Christian Kroll, Bilhan Karacora, Sven G. Meuth, Gerhard Fritz, Orhan Aktas, Carsten Berndt, Tim Prozorovski

## Abstract

DNA damage is a major risk factor for the decline of neuronal functions with age and neurodegenerative diseases. The connection between DNA damage and neurodegeneration is extensively investigated, however, the mechanisms limiting the propagation of damaged DNA from highly replicative neural stem/progenitor cells (NSPCs) to post-mitotic neurons remains largely unknown. Here, we describe that enzymatic activity of the histone deacetylase sirtuin 1 (SIRT1) is important for the homologues recombination-dependent repair of double-stranded DNA breaks induced by etoposide. Furthemore, SIRT1 abolishes neuronal fate of murine NSPCs following induction of DNA damage. Pharmacological inhibition or genetic inactivation of SIRT1 rescues etoposide-mediated inhibition of neuronal differentiation in NSPCs and hippocampal slice cultures and promotes transcription of pro-neuronal genes. Inhibition of Ataxia-telangiectasia mutated (ATM), the central regulator of the DNA damage response, mimics the SIRT-dependent effect of DNA damage on neuronal differentiation, indicating that the ATM/SIRT1 axis inhibits formation of neuronal cells harbouring damaged DNA. These data are consistent with the role of SIRT1 in genome stability, healthy ageing and protection from the development of ageing-associated neurodegenerative diseases.

## Introduction

In *Drosophila* larval brain, induced DNA damage in NSPCs (called neuroblasts) rapidly induces cell cycle exit and subsequent premature neuronal differentiation (Wagle & Song, 2020; Xu et al., 2023). This mechanism allows to withdraw the cells with affected genome from the pool of proliferating precursor cells and thus, to minimize propagation of mutations and genomic aberrations (Jackson & Bartek, 2009). By keeping this mechanism (Barazzuol et al., 2017), mammals additionally solve this problem by developing complex DNA damage repair mechanisms or by triggering cell death (apoptosis) or cellular senescence in cells burdened by severely damaged DNA (Ribeiro et al., 2023). Nevertheless, increasing evidence indicates that DNA damage occuring due to insufficient DNA repair may accumulate in post-mitotic neurons and is linked to altered neuronal function (Konopka & Atkin, 2022) and neurodegeneration (Homma et al., 2021; Delint-Ramirez & Madabhushi, 2025). Thus, the knowledge of neurobiological mechanisms that limits the spreading of damaged DNA in neural tissue, e.g. by supporting mechanisms involved in DNA repair or withdrawal of injured cells, is of therapeutic interest, particularly for neurological complications of chemotherapy (Chemobrain; chemotherapy-induced cognitive impairment) (Torre et al., 2021; Murillo et al., 2023). Here, we eximined the neuronal fate of murine neuronal stem/progenitor cells (NSPCs) after induction of DNA double strand breaks (DSBs). We focused on the NAD+-dependent histone deacetylase sirtuin 1 (SIRT1, HDAC class III), whose function is implicated in the maintenance of genome integrity (Oberdoerffer et al., 2008; Wang et al., 2008), neural stem cell fate (Prozorovski et al., 2015) and serves as a therapeutic target in neurodegeneration (Thapa et al., 2024), chemotherapy (Cheng et al., 2003) and healthy ageing (Herranz et al., 2010; Satoh et al., 2013).

We hypothesize, that SIRT1 may be an attractive candidate to link DNA damage response in NSPCs to cell fate determination. This hypothesis was also strengthened by previous observations of the critical role of SIRT1 in suppression of neuronal commitmence upon mild oxidative stress (Prozorovski et al., 2008) or mitochondrial DNA damage (Wang et al., 2011). Here, using cellular and hippocampal slice cultures with loss of SIRT1 activity either by using genetically inactivated SIRT1 mice (Cheng et al., 2003) or the clinically relevant pharmacological SIRT1 inhibitor Ex-527/Selisistat and treatment with low dosage of the topoisomerase II inhibitor etoposide we show that 1) SIRT1 promotes homologous recombination (HR)-mediated DNA repair in mitotically active NSPCs, 2) NSPCs deficient for SIRT1 exhibit elevated levels of DSB, and 3) DNA damage abrogates neuronal fate unless cells lack SIRT1 activity.

Taken together, our data emphasize SIRT1 as an important factor for genomic integrity of neural tissue and highlight the need for cautious use of SIRT1 inhibitors.

## Material and Methods

### Animals

Animal handling complied with German animal protection law (TierSchG) and received approval from Landesamt für Natur, Umwelt und Verbraucherschutz Nordrhein-Westfalen (LANUV) (O74/08). Experiments utilized SIRT1 mice lacking exon 4 (ΔSIRT1) generated by Cheng et al., 2003 (Cheng et al., 2003) and control SIRT1-expressing littermates under C57BL/6 background. ΔSIRT1 mice express a catalytically inactive mutant SIRT1 form specifically in nestin-expressing cells. Genotyping involved cerebellum DNA isolation and PCR analysis using specific primers.

### Isolation and cultivation of murine NSPCs

NSPCs from dissected subventricular zones (SVZ) and the hippocampi were isolated from postnatal day 2-5 mouse pups by pooling cells from a minimum of three animals. Isolation followed protocol by Guo et al., 2012 . Briefly, mice were decapitated, brains sectioned into 800 µm coronal slices, and lateral SVZ and hippocampi collected. For tissue dissociation Miltenyi’s adult brain dissociation kit (Miltenyi, 130-107-677) was used. After passing through a 70 µm mesh, cells were washed twice in HBSS (Cat.: 14025092; Thermo Fisher Scientific). NSPCs were expanded as free-floating neurospheres in neural proliferation media (NPM; Neurobasal media (Cat.: 21103049; Thermo Fisher Scientific), 1% penicillin-streptomycin (Cat.: 15070063), 2 mM Glutamax (Cat.: 35050061; Thermo Fisher Scientific), 2% B27 (without retinoic acid) (Cat.: 12587001; Thermo Fisher Scientific), 20 ng/ml bFGF and EGF (growth factors; Cat.: 12343407 and 12343627, respectively; Immunotools GmbH) at 37°C, 5% CO2 (incubation). NSPCs were sub-cultured every 4 days up to 3-5 passages on poly-L-ornithine (PLO)-coated flasks in NPM without compromised proliferation. Cultivating media was screened for mycoplasma infection. The following agents all dissolved in DMSO were used for treatment: Ex-527 (5 µM; Cat.:2780, Tocris), SRT 1720 (Cat.: 7558; Tocris), Ku- 55933 (10 µM; Cat.: Cat. No. 3544; Tocris), SCR7 (10 µM; Cat.: 74102; Stem Cell Technology), B02 (10 µM; Cat.: 6392; Tocris)

### Differentiation of NSPCs

NSPCs were seeded on PLO/Laminin-coated glass cover slips. Differentiation was induced by growth factor withdrawal, using neuronal differentiation media (NDM; Neurobasal media, 1% P/S, 2 mM Glutamax, 2% B27 (with retinoic acid; Cat.: 17504044; Gibco), and 1% N2 (Cat.: 17502048; Gibco). Half of the media was exchanged with fresh NDM every second day.

### Cell viability assay

Cell viability was assessed using the Resazurin reduction assay (CellTiter-Blue®; Promega, G8081). Sub-cultured NSPCs (1 x 10^4^ cells in 100 µl NPM) were incubated for 2 days in poly-L-lysine (PLL)-precoated 96-well plates (Corning). Six technical replicates per group were averaged. A final concentration of 0.01% Triton-X 100 added to the media served as positive control. CellTiter-Blue® reagent (1:6) was applied, and fluorescence was measured using a Magellan plate reader.

### Immunocytochemistry

Cell cultures grown on PLO-coated 13 mm glass cover slips in 24-well plates were fixed with 4% paraformaldehyde (ROTI^®^Histofix; Cat.: P087.1; Carl Roth) for 15 min at room temperature (RT, 20°C), washed in PBS and blocked with 5 % goat serum (Cat.: 10189722; Fisher Scientific) in 0.25% Triton X-100/PBS for two hours at RT. Primary antibodies were diluted in blocking buffer and incubations were performed overnight at 4°C (primary antibody). The following antibody were used: mouse anti-H2A.X (Ser139) (Cat.: 50-9865-42; eBioscience; 1:1000); rat anti-53BP1 (Cat.: 933001; Biolegend; 1:500); rabbit anti-RAD51 (Cat.: 133534; Abcam; 1:1000). Bound antibodies were visualised after one hour incubation at RT by using appropriate combinations of species/isotype-specific fluorochrome-conjugated secondary antibodies: goat anti-rabbit IgG (Cy3, Merck Millipore, 1:500), goat anti-rabbit IgG (Alexa Fluor 488, Jackson ImmunoResearch, 1:500), goat anti-rat IgG (Cy2, Merck Millipore, 1:500). Nuclei were counterstained with Hoechst 33258 (2 µg/ml; Invitrogen;) for 5 minutes. Coverslips were mounted on glass slides in Immu-mount^®^ (Fisher Scientific, 10622689) medium. Coverslips were imaged using an Olympus BX51 fluorescence microscope. All Images were contrast-enhanced in a comparable fashion using Adobe Photoshop software to facilitate visibility in composite figures.

### γH2AX foci analysis

At least 50 nuclei from 4 fields were assessed at 100 x magnification. To automatically count foci, images were processed and analyzed with ImageJ softaware (FIJI; https://imagej.nih.htmlgov/ij/index, v2.0.0-rc-69/1.52n). To outline nuclei-ROI, image filtering, thresholding, and watershed segmentation was used. γH2AX images were augmented, and foci converted to single pixel. The nuclei-ROI mask was then used to count pixels per nucleus.

### Western blotting

Cells were scraped into cold DPBS on ice, pelleted with centrifugation (200 x g; 5 min; 4°C), and washed with cold DPBS. Pellets were lysed in RIPA buffer (Thermo Scientific, 89900) supplemented with protease and phosphatase inhibitors (Roche, 11873580001 or Thermo Scientific, A32957) for 10 min on ice. After sonication and centrifugation (16 000 x g; 10 min; 4°C), lysate protein concentration was measured with BCA kit and samples were set to 1,5 µg / µl using a BCA kit. Lysates were mixed with 4x Laemmli sample buffer (BioRad; 1610747) supplemented with reducing buffer. 25-50 µg of proteins were loaded for analysis. Gel electrophoresis and blotting procedure was done using BioRad Any kD™ Mini-PROTEAN® TGX Stain-Free™ Protein Gels (BioRad; 4568123) and commercial BioRad equipment and buffer. Membranes were blocked with EveryBlot Blocking Buffer (BioRad; 12010020) or 5% BSA in PBS containing 0.05 % Tween-20 (TBS buffer). Primary and secondary antibodies were prepared in 2.5% BSA in TBS and incubated for 2h at RT or overnight at 4°C with shaking, followed by washing with TBS. Stain-free development and imaging, as well as infrared imaging of immuno-stained membranes, were conducted using ChemiDoc Imaging Systems (BioRad) detecting at 680 and 800 nm. Densitometry analysis was performed using BioRad’s ImageLab software and ImageJ. For quantitative analysis, protein expression was normilized to the expression levels of the houskeeping protein β-actin. The following primary antibody were used: mouse anti-H2A.X phospho-Ser139 (Cat: 613402; Biolegend; 1:1000), rabbit anti-acetyl-p53 (Lys320) (Cat.: 06-1283; Millipore; 1:1000), rabbit anti-Sirt1 (Cat.: 07-131; Millipore; 1:1000), rabbit anti-RAD51 (Cat.: 133534; Abcam; 1:1000), mouse anti-β-actin (Cat.: A5316; Sigma-Aldrich; 1:5000)

### Redox shift analysis

*In situ* thiol modifications of SIRT1 were assessed using indirect redox shift by using electrophoretic mobility shift assay (Habich & Riemer, 2017) . NSPCs (2 x 10^5^ cells) were cultivated on PLO-coated 6-well plates for 2 days. After treatment, cells were incubated with ice-cold 20 mM N-ethylmaleimide (NEM; Cat.: 128-53-0; Merck) in PBS for 10 minfollwed by lysation in 8% trichloroacetic acid (TCA). The precipitates were collected and centrifuged (16 000 x g; 10 min; 4°C), washed with 5% TCA, and resuspended in 2x Laemmli buffer (BioRad, 1610737) with 20 µM reducing agent tris(2-carboxyethyl)phosphine (TCEP; Cat.: C4706, Sigma-Aldrich). pH was adjusted with 1 M Tris-HCl to pH 8.5. Samples were sonicated until the pellet was dissolved. Thiol-reactive probes (NEM or mPEG24 (Sigma-Aldrich)) at 20 mM were added, and samples were kept in the dark at RT for 1h. Western blot analysis with SIRT1 immuno-detection was performed as described above. Reaction of reduced thiols with mPEG24 causes protein band shifting in gel electrophoresis, correlating to the number of reduced thiols present.

### RNA isolation and quantitative PCR analysis

RNA isolation from cultured cells was performed using TRIZOL^®^ solution according to the manufactural’s instructions (Invitrogen). Purity and amount of RNA were measured with Nanodrop 2000 (Thermo Scientific). cDNA was synthesized with Taqman high capacity reverse transcription reagents (Applied Biosystems) from 2 µg DNA ; qRT-PCR was performed with Power SYBRGreen fluorescent dye (Applied Biosystems) and the 7000 Real Time PCR System (Applied Biosystems). Primers that span an exon-exon junction were designed using Primer Express software (Applied Biosystems). Each measurement was performed in duplicates. For all samples, *glyceraldehyde 3-phosphate dyhydrogenase* (*GAPDH*) was used as housekeeping gene.

### Hippocampal Organotypic Slice Culture (hOSC)

hOSCs served as an *ex vivo* model for investigating neurogenesis in a physiological tissue setting (Raineteau et al., 2004). Slices were prepared from postnatal day 3 (P3) mice and cultured following a modified standard interface method as described previously (Schneider et al., 2016). Briefly, brains from decapitated mice were obtained and hippocampi isolated. The hippocampi were sliced into 400 µm thick coronal sections using a McIllwain tissue chopper (GaLa Instrumente). Only intact slices from the hippocampus, excluding regions near spatial and temporal poles, were selected. These slices were placed on porous membrane inserts (Millicell-CM PICM03050; Millipore) in 1 ml warm and CO2-saturated slice culture medium (50% MEM containing 2 mM glutamax (Cat.; 32561029), 25% heat-inactivated horse serum (Cat.: 16050122), 25% HBSS (Cat.: 14025092), 1% penicillin-streptomycin (P/S; Cat.: 15070063), 6.5 mg / ml D-glucose (Cat.: A2494001); all from Thermo Fisher Scientific) in 6- well plates. After 24h incubation, slice medium was replaced for serum-free hippocampal organotypic differentiation media (HDM; Neurobasal media, 1% P/S, 2 mM Glutamax, 2% B27 (with retinoic acid; Cat.: 11530536; Fisher Scientific) with or without treatment, and hOSCs were incubated for an additional 24h before daily media exchange. Slices were analysed 8 days after etoposide treatment. To fixate hOSCs, inserts were washed and submerged in 4% PFA. Slices were stored in PBS. For immuno-staining, the membrane was permeabilized in 0.5% Triton X-100 in PBS. The membrane was mounted on slides using Immu-mount®, dried overnight, and imaged by Zeiss confocal microscopy with complete z-stack and merging using mosaic-merge technique.

### Statistical analysis

Statistical analysis was performed using Student’s t test or one-way ANOVA followed by Tukey’s post hoc test. Data are represented as mean ± standard error of mean (SEM). In all experiments a p value: *<0.05; **<0.01; ***<0.001 was defined as a statistically significant. All statistical analyses were performed using GraphPad Prism 5 software (RRID:SCR_015807).

## Results

### Loss of SIRT1 activity does not sensitize NSPCs to etoposide

Taking into account the multiple regulatory roles of SIRT1 in DDR and genome stability, we first examined the involvement of SIRT1’s enzymatic activity in DSB repair in cultured NSPCs. Mouse NSPCs were isolated from postnatal brains of Nestin^Cre(+)^SIRT1^LoxP(ex4)^ (hereafter ΔSIRT1; Cheng et al., 2003) and corresponding littermates Nestin^Cre(-)^SIRT^LoxP(ex4)^ (hereafter control mice).

Mixed population of the SVZ- and SGZ-derived NSPCs were cultivated in proliferating conditions that support growth of EGF and bFGF-responsive multipotent and intermediate progenitors (Lim & Alvarez-Buylla, 2016; Song et al., 2012). After the addition of increasing dosages of the topoisomerase II inhibitor etoposide/VP-16, cell viability was not significantly different between ΔSIRT1 and control NSPCs (Fig. 1A).

**Fig. 1.**
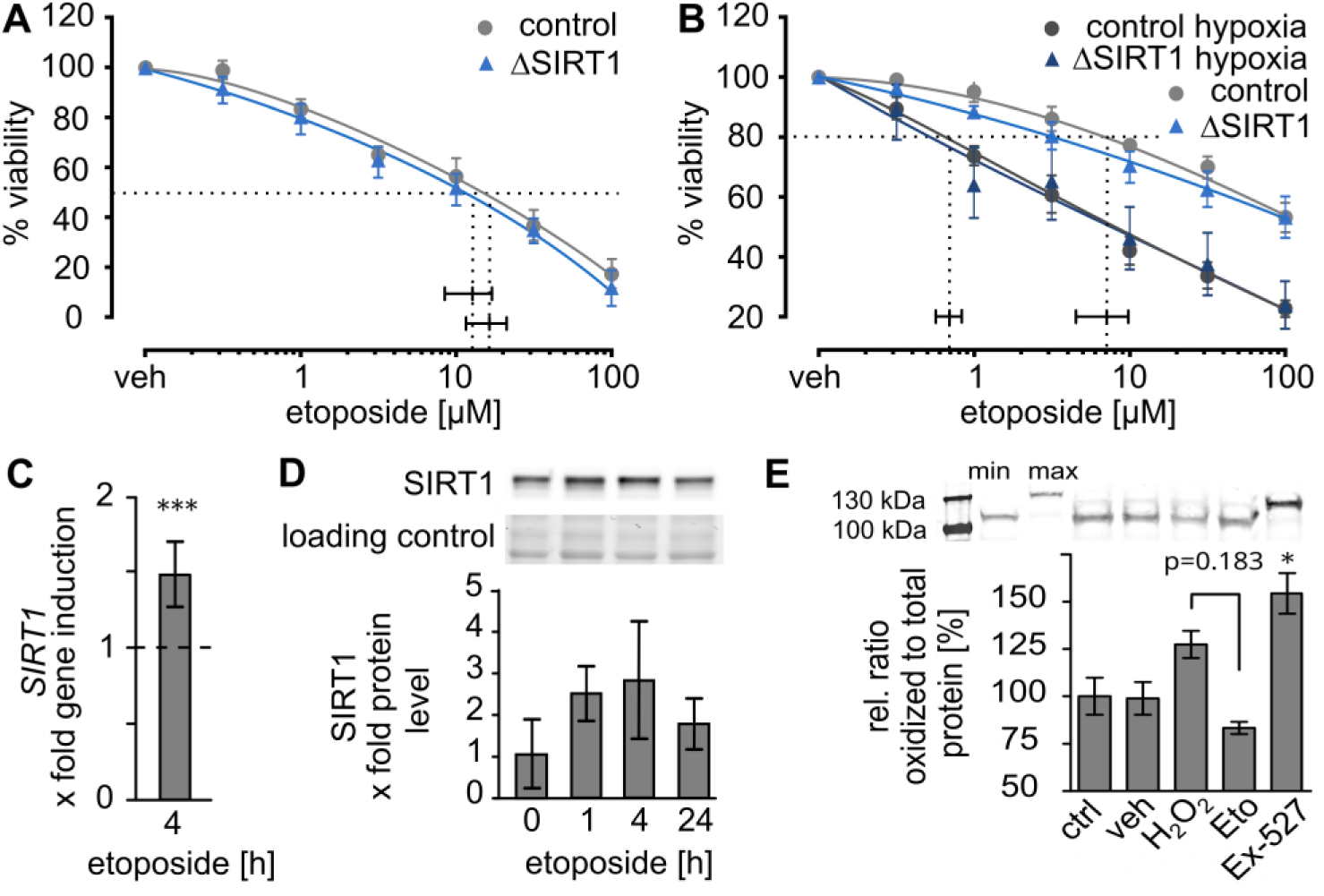
SIRT1 activity is increased upon etoposide treatment but does not affect survival of NSPCs. **A, B)** NSPCs isolated from control and ΔSIRT1 mice were treated for 24 h (**A**) or for 1 h following 23 h of recovery with or without hypoxia (5 % O2) (**B**) with indicated concentrations of etoposide. Cell survival was measured by CTB assay (n=3). **C, D)** Treatment with 1 µM etoposide for indicated time points increased transcription **(C,** n=3) and translation of SIRT1 **(D,** n=3). **E)** Treatment with 1 µM etoposide for 1 h leads to active, reduced SIRT1 (in contrast to H_2_O_2_ and Ex-527 treatment) as shown by redox shift western blots (n=3). Significance was calculated using one-way ANOVA followed by Tukey’s post hoc test or unpaired two-tailed Student’s t-test, as appropriate, *: p<0.05, ***: p<0.001.

Similarly, we did not observe differences in cell survival in control NSPCs treated either with SIRT1 inhibitor (Ex-527; Napper et al., 2005; Solomon et al., 2006), or SIRT1 activator (SRT1720; Milne et al., 2007) in addition to etoposide (Supplementary Fig. S1A). We also tested the sensitivity towards etoposide under mild hypoxic condition that mimics lowered oxygen levels in germinative zones of the brain (De Filippis & Delia, 2011). Consistent with previous findings (Ng et al., 2018), treatment of NSPCs cultivated at 5% O_2_ increased sensitivity to etoposide. Nevertheless, similar to normoxic conditions, there were neither significant differences between control and ΔSIRT1 cultures (Fig. 1B), nor in control NSPC cultures with modulated SIRT1 activity (Supplementary Fig. S1B). SIRT1 itself is upregulated upon treatment with etoposide on RNA and protein levels (Fig. 1C, D). Moreover, etoposide treatment leads to the formation of the active, reduced form of SIRT1, whereas Ex-527 drastically diminished the active form of SIRT1 (Fig. 1E). Numbers of nuclear γH2AX foci (DSB marker, Mah et al., 2010) significantly increased in etoposide-treated ΔSIRT NSPCs compared to controls (Fig. 2A). Especially repair of DSBs seems to be affected in ΔSIRT NSPCs or NSPCs treated with Ex-527 as shown by less efficient decreased γH2AX number (Fig. 2A).

**Fig. 2.**
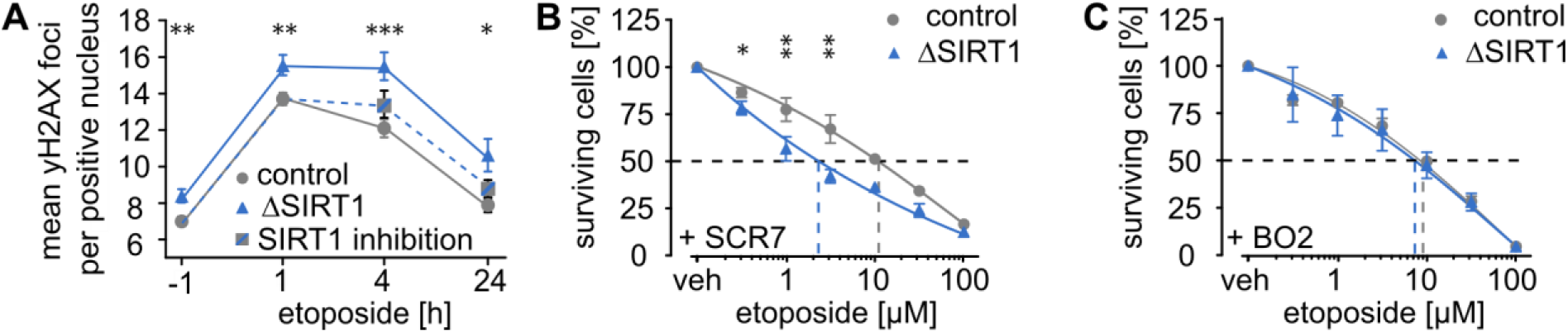
SIRT1 supports DNA damage repair via homologues recombination. **A)** After treatment of NSPCs with 1 µM etoposide for 1 h, γH2AX foci were counted at different time points during recovery in control cells and cells lacking SIRT1 activity (n=3; ≥50 cells per replicate). NSPCs isolated from control and ΔSIRT1 mice were treated for 24 h with indicated concentrations with or without presence of 50 µM SCR7 **(B)** or 50 µM BO2 **(C)** and survival was determined using the CTB assay (n=3). Significance was calculated using two-way ANOVA followed by Tukey’s post hoc test, *: p<0.05, **: p<0.01, ***: p<0.001.

Exposure to low etoposide concentration leads to clear puncta of γH2AX typical for DNA DSBs, rather than pan-nuclear staining typical for apoptosis (Shimada et al., 2015). Following one hour of exposure to etoposide, NSPCs were released into DNA repair phase by re-incubation in fresh proliferation medium. Quantitative analysis of γH2AX foci showed a decline of γH2AX foci within the first four hours and after 24 hours γH2AX signals returned to basal level in control cultures, reflecting DNA repair. In ΔSIRT1 cells elevated numbers of γH2AX foci were evident in NSPCs prior and following treatment with etoposide (Fig. 2A). In contrast to control cells, high number of γH2AX foci was stable after four hours in ΔSIRT1 and those treated with Ex- 527 (Fig. 2A) and persisted after 24 hours..

### Role of SIRT1 in homologues recombination and non-homologues end joining

Our data indicate that NSPCs deficient for SIRT1 activity exhibit impaired DNA damage repair. Considering the competitive nature of the two main DSB repair mechanisms — HR and non-homologous end joining (NHEJ) — we tested the potential compensatory involvement of each pathway in ΔSIRT1 cells. Inhibition of NHEJ-mediated DNA repair via DNA ligase IV inhibition (SCR7 inhibitor) significantly sensitized ΔSIRT1 NSPCs to etoposide-induced DNA damage as compared to control cells (Fig. 2B). Treatment with BO2, a selective HR inhibitor, had no additional detrimental effect on cells lacking SIRT1 activity (Fig. 2C). These results point to the reliance of SIRT1 deficient cells on the NHEJ repair indicating that SIRT1 is especially important for HR-mediated DSB repair in NSPCs. In addition, we did not observe a difference in the number of γH2AX^+^/53BP1^+^ foci between control and ΔSIRT1 cells (Fig. 3A). Since 53BP1 is important for early response to DSBs and a main regulator of NHEJ (Lei et al., 2022), this supports the finding that NHEJ is not affected by the loss of SIRT1 activity (Fig. 3A, B). Next, we tested whether SIRT1 impairment impacts HR regulation. The number of γH2AX foci co- localized with the recombinase RAD51 was significantly decreased in ΔSIRT1 NSPCs (Fig. 3A, C). RAD51 catalyzes the core reactions of HR in the late phase of DSB repair (Krishna et al., 2007), and the lack ofγH2AX^+^/RAD51^+^ foci indicates that ΔSIRT1 NSPCs have lower capacity for HR as in control cells RAD51 foci formation is regulated by increased RAD51 expression or protein stabilization via post-translational modifications (Orhan et al., 2021).

**Fig. 3.**
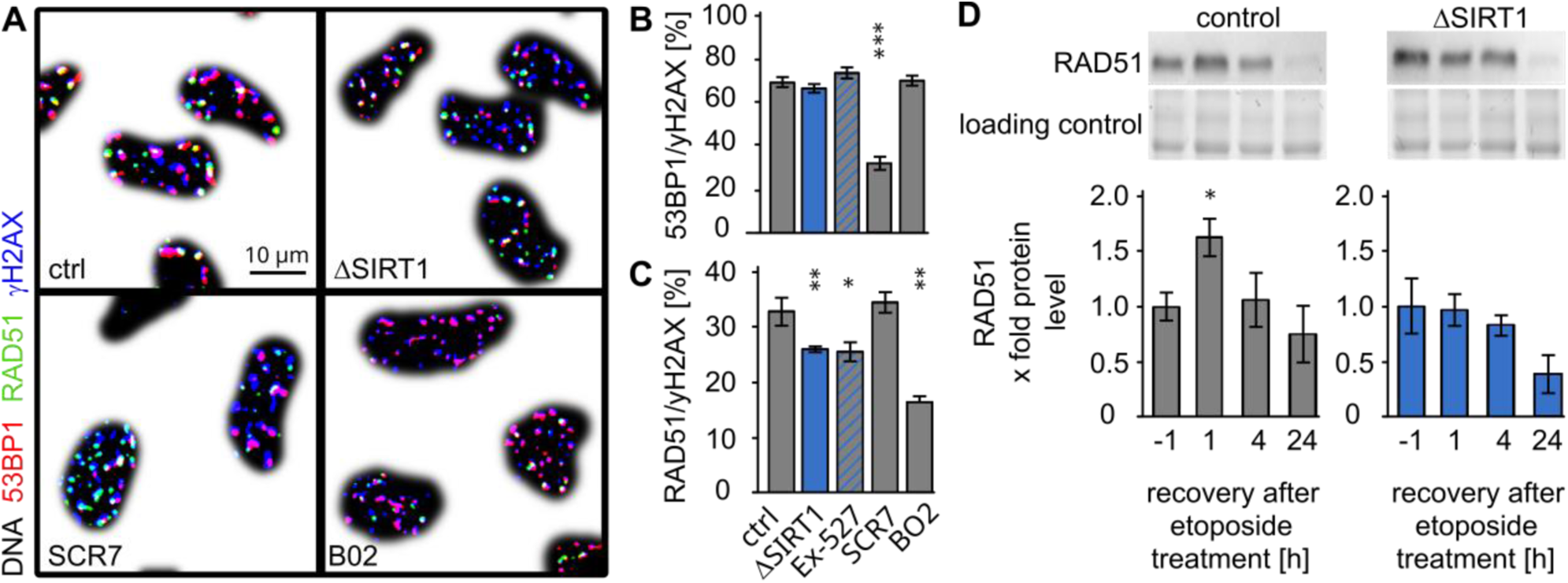
Lack of SIRT1 activity interferes with RAD51-dependent homologues recombination. After immunocytochemistry using anti-RAD51-, -53BP1-, and -γH2AX- antibodies in combination with Hoechst **(A)**, co-localization of γH2AX foci with RAD51 or 53BP1 in the nucleus of control cells, ΔSIRT1 cells and cells treated with Ex-527, SCR7, or BO2 were counted 4 h after etoposide insult (1µM, 1 h) **(B, C,** n=3). **D)** Protein levels of RAD51 after different time periods of recovery after treatment with 1 µM etoposide were determined in control and ΔSIRT1 NSPCs by western blots (n=3). Significance was calculated using two-way ANOVA followed by Tukey’s post hoc test, *: p<0.05, **: p<0.01, ***: p<0.001.

Induction of RAD51 in response to DNA damage was abolished in ΔSIRT1 NSPCs (Fig. 3D) and cells treated with Ex-527 (Fig. S2A). Expression and activity of RAD51 upon DNA damage is affected among others by p53 (Bertrand et al., 2004; Arias-Lopez et al., 2006). In this regard, we observed that SIRT1 inhibition upregulates p53 acetylated at lysine K320 (Fig. S2B), a post-translational modification that stabilizes the protein (Knights et al., 2006).

### Etoposide-mediated abrogation of neuronal generation requires SIRT1 activity

After three and seven days of differentiation, only a minor population of NSPCs adopt neuronal phenotype and express the pan-neuronal marker βIII-tubulin when differentiation was induced four hours after recovery from etoposide bolus treatment (Fig. 4A, B). This diminished ability to generate neurons was not observed in NSPCs lacking SIRT1 activity; almost the same amount of neuronal cells was formed with and without etoposide (Fig. 4A, B). This NSPC response to DNA damage was also observed in hippocampal slice cultures from age-matched postnatal ΔSIRT1 and control mice. Here, we used doublecortin, an early neural differentiation marker (Gleeson et al., 1998) to follow neuronal fate. Again, etoposide treatment leads to fewer neurons, whereas in slice cultures obtained from ΔSIRT1 this effect was lost (Fig. 4C, D). This effect goes along with increased transcription of pro-neuronal genes *Mash1*, *Math1*, *Pax3*, and *Prox1* at early stages of differentiation upon Ex-527 treatment (Fig. 4E). These cells were exposed to etoposide for one hour and after four hours of recovery, differentiation was induced for 20 h. At the same time point, γH2AX levels were increased in the cells with SIRT1 inhibition compared to control cultures (Fig. 4F, G).

**Fig. 4.**
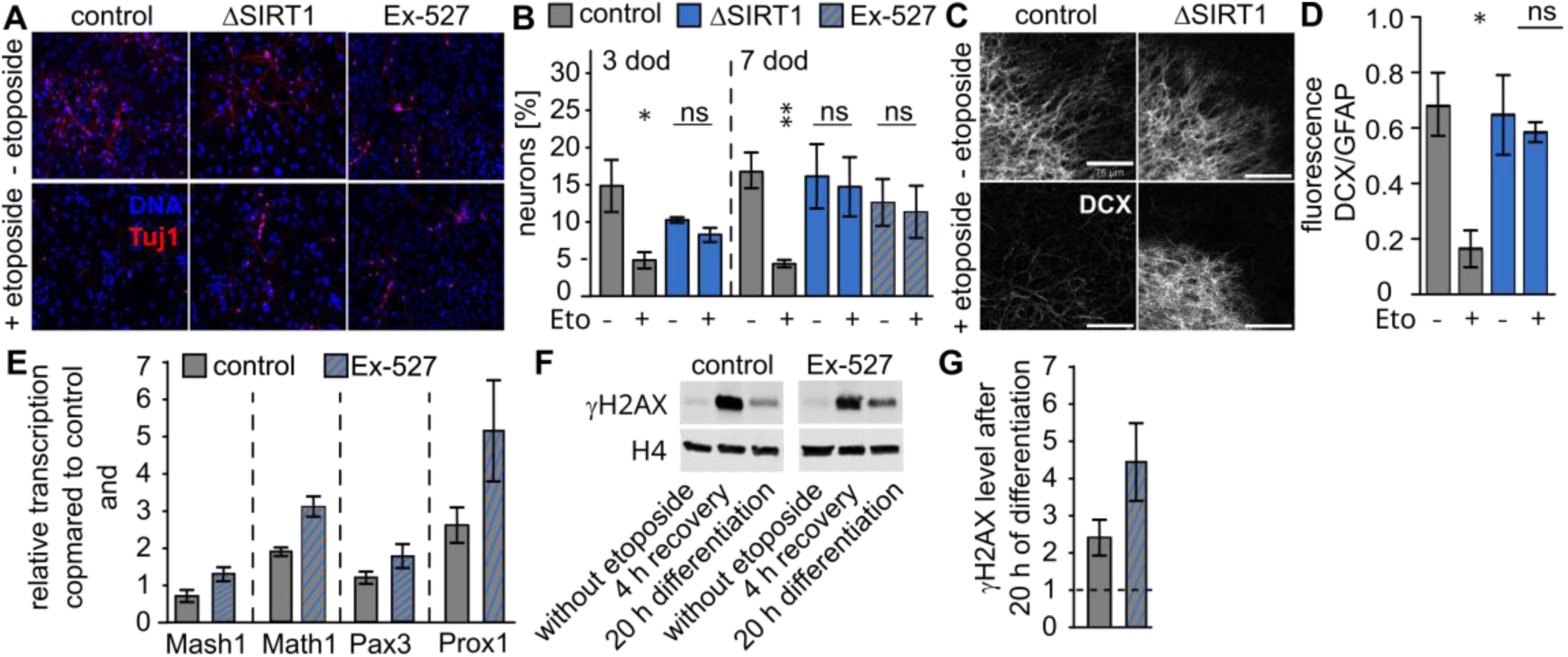
Deficiency of SIRT1 activity allows differentiation towards neuronal progeny with increased amounts of DNA damage. **A)** Tuj1 positive neurons were visualized by respective antibodies after seven days of differentiation of NSPCs isolated from control mice (± Ex-527) and ΔSIRT1 mice. Before differentiation, cells were treated with or without 1 µM of etoposide for 1h and allowed to recover for 4 h. **B)** shows quantification of neuronal progeny after three and seven days of differentiation (n=3). **C)** Doublecortin (DCX) positive neurons were stained via respective antibodies in hippocampal organotypic slice cultures isolated from control or ΔSIRT1 mice ± etoposide treatment (1 µM for 24 hours) after 10 days of differentiation (scale bar = 75 µm). **D)** shows quantification of DCX positive neurons (n=3). **E)** NSPCs were treated with 1 µM etoposide for 1 h with or without presence of Ex-527. After four hours of recovery, differentiation was induced for 20 h. At this time point mRNA was isolated and transcription of the mentioned genes was determined by qRTPCR (n=3). **F)** In lysates of cells treated as described under **E)** γH2AX levels were visualized by western blots. **G)** Quantification of **F)** (n=3). Significance was calculated using two-way ANOVA followed by Tukey’s post hoc test, *: p<0.05, **: p<0.01.

### ATM/SIRT1 axis regulates NSPĆs fate after DNA damage

ATM is the central regulator of the DNA damage response and phosphorylates a broad spectrum of substrates that are virtually linked to NSPC functions. Therefore, we tested whether ATM activity is required for the effects described above. NSPC differentiation showed that ATM works in the same line as SIRT1. Inhibition of ATM (treatment with Ku-55933) rescues diminished differentiation of NSPCs into βIII-tubulin cells upon etoposide treatment (Fig. 5A, B). Inhibition of ATM antagonized the effect of etoposide on transcription of the same pro-neuronal genes (Fig. 5C) as shown for SIRT1 inhibition.

**Fig. 5.**
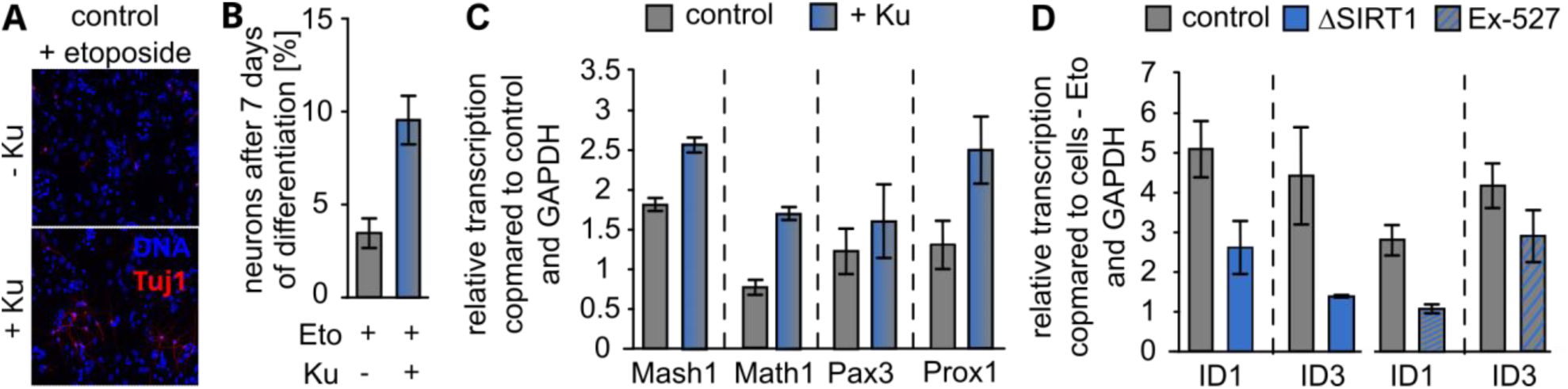
SIRT1 acts in line with ATM in DNA damage recovery. **A)** Tuj1 positive neurons were visualized by respective antibodies after seven days of differentiation of NSPCs isolated from control mice ± Ku-55933 (Ku; ATM inhibition). **B)** shows quantification of A) (n=3). **C)** mRNA was isolated from NSPCs differentiated for seven days ± Ku and transcription of indicated pro-neuronal genes was determined by qRTPCR (n=4). **D)** Transcription levels of ID1 and ID3 measured by qRTPCR in ΔSIRT1 and control NSPCs (± Ex-527) after 1h of treatment with 1 µM etoposide, 4 h recovery, and 20 h of differentiation (n=3).

In response to DNA damage, we also observed induction of downstream targets of the ATM activated BMP/SMAD signalling pathway (Chau et al., 2012), Id1 and Id3 genes (Fig. 5D). Our data revealed that transcription of *Id1* and *Id3* genes was abolished by SIRT1 inhibition suggesting that SIRT1-dependent allowance for neuronal differentiation upon DNA damage is facilitated in concert with the ATM/SMAD axis.

## Discussion

Growing evidence indicates that DNA damage response is interconnected to developmental (Wei et al., 2016; Alt & Schwer, 2018; Qing et al., 2023; Alt et al., 2017) and metabolic signalling pathways (Kwon et al., 2019; Moretton & Loizou, 2020). However, factors that link these processes remain incompletely understood. In this work we focused on the epigenetic modulator SIRT1, a NAD+-dependent deacetylase, that has been previously implicated in the regulation of metabolism (Chang & Guarente, 2014), cancer cell reprogramming (Filipponi et al., 2019), cell fate decision of neural progenitors (Prozorovski et al., 2008; Prozorovski et al., 2019), lineage commitment following DNA damage (Rimmelé et al., 2014), and DNA repair (Lagunas-Rangel, 2019). We demonstrated that SIRT1 inhibits neuronal fate of NSPCs after DNA damage. We propose that this mechanism inhibiting accumulation of DNA damage protect against neurodegeneration.

### Lack of SIRT1 activity allows neuronal differentiation of NSPCs with damaged DNA

Our main finding is that SIRT1 negatively regulates neuronal fate upon DNA injury, whereas loss of SIRT1 activity permits differentiation of NSPCs harbouring elevated levels of DNA damage. Such failure to repair DNA lesions properly after the induction of cell proliferation arrest can lead to mutations or large-scale genomic instability, hallmarks of tumorigenicity and neurodegeneration (Qing et al., 2023). Such scenario was recently observed in mice expressing a mutated form of VCP. This protein is involved in DNA damage repair in NSPCs and linked to frontotemporal lobal degeneration. Accumulated DNA damage in mutant NSPCs led to delayed neurodegeneration in adult mice (Homma et al., 2021). Mice with genetic inactivation of SIRT1 in NSPCs have no overt effect on neural development. However, aged SIRT1-deficient mice have been reported to show increased accumulation of DSB in SVZ NPCs (Ren et al., 2022), and cultured post-mitotic neurons deficient for SIRT1 activity showed increased sensitivity to DNA damage (Dobbin et al., 2013). Following etoposide treatment SIRT1 transcription and protein level was transiently upregulated in NSPCs. We also observed increase in its reduced, active state (Caito et al., 2010; Bräutigam et al., 2013) confirming the need of its activity during DNA damage repair.

### SIRT1 acts in concert with ATM and promotes homologues recombination

DNA damage repair is facilitated by HR or NHEJ. Both mechanisms are regulated by ATM, a kinase with many substrates. Important for this study is its function in elimination of neural cells that display genomic damage (Lee et al., 2000; Shull et al., 2009). We have shown that pharmacological ATM inhibition mimics the loss of SIRT1 activity after etoposide treatment (increased transcription of pro-neuronal genes and number of neurons) and that ATM- controlled transcription of *Id1*and *Id3*, two SMAD-dependent genes encoding proteins interfering with neuronal differentiation, are less abundant in NSPCs with diminished SIRT1 activity. The above mentioned results indicate that SIRT1 and ATM works in the same direction during DNA damage response. Indeed, interaction between SIRT1 and ATM has been shown before. Moreover, activation of SIRT1 diminished the phenotypes caused by ATM deficiency (Fang et al., 2016).

Based on our data we conclude that SIRT1 is more important for HR than for NHEJ in mitotic cells. Consistently, cells lacking active SIRT1 are more susceptible towards NHEJ inhibition over HR suggesting that NHEJ is the predominant repair pathway in ΔSIRT1 NSPCs or NSPCs treated with Ex-527. We did not observe any effect of SIRT1 deficiency on the number of γH2AX foci co-localized with a marker protein of NHEJ, 53BP1. However, the number of γH2AX/Rad51 (a marker protein of HR) foci were diminished in cells lacking SIRT1 activity in agreement with previous research (Wang et al., 2008; Rasti et al., 2023; Oberdoerffer et al., 2008). Moreover, SIRT1 inactivation led to inability of rising RAD51 protein level upon damage, the mechanism that may potentially involve accumulation of acetylated p53, an inhibitor of Rad51 expression and HR (Bertrand et al., 2004; Arias-Lopez et al., 2006). SIRT1 is a well-known inhibitor of p53 by promoting p53 degradation and its inactivation sensitizes cancer cells to DNA damaging agents such as etoposide (Hajji et al., 2010; Zhang et al., 2016; Kweon et al., 2014). Suppression of p53 activity by SIRT1 has also been implicated in survival and maintenance of neural cancer stem cells (Lee et al., 2015). In conclusion, our data suggest a novel mechanism by which SIRT1-p53 interplay mediates genomic stability in non-cancer cells and broadens its regulatory role in the HR-dependent DNA repair. However, in post-mitotic neurons, SIRT1 is required for efficient DSB repair via NHEJ (Dobbin et al., 2013).

### SIRT1 inhibition promotes neuronal fate under physiological and pathological conditions

The ability of SIRT1 to modulate NSPC fate has been demonstrated in *in vitro* and *in vivo* experiments. We have shown before that SIRT1-deficient embryonic NSPCs are resistant to inhibition of neurogenesis under mild oxidative stress (Prozorovski et al., 2008). Loss of SIRT in NSPCs promotes neuronal differentiation (Rafalski et al., 2013), whereas SIRT1 overexpression blocks neuronal differentiation *in vitro* and in the CNS germinative zones (Liu et al., 2014; Saharan et al., 2013).

### Is SIRT1’s function in DNA damage response in NSPCs connected to protection against neurodegeneration?

We hypothesize that our data showing that SIRT1 activity inhibits formation of neurons from NSPCs harbouring damaged DNA might be a relevant mechanism for protection against neurodegeneration. This assumption is consistent with data demonstrating hippocampal atrophy and cognitive impairment in mice with conditional SIRT1 knockdown (Sun et al., 2022). Increased SIRT1 function through overexpression or the use of SIRT1 activators enhance neuronal survival (Guo et al., 2011). Whether all these findings are connected to DNA damage repair or SIRT1-dependent decision on the DNA damage response remains to be investigated. However, modulation of SIRT1 is considered a novel therapeutic strategy in various neurodegenerative diseases (Mishra et al., 2021)

## Author contributions statement

Conceptualization and supervision: OA, CB, TP; critical input: SGM, GF, OA; data generation: CK, BK,TP; data analysis: CK, CB, TP; manuscript preparation: CB, TP; figure preparation: CK, CB, TP. All authors have read, revised and approved the final manuscript.

## Data Availability Statement

Data supporting the findings of this study are available within the figures and the results section. Further data and extended description of methods are available from the corresponding authors upon reasonable request.

## Supplementary information

**Fig. S1.**
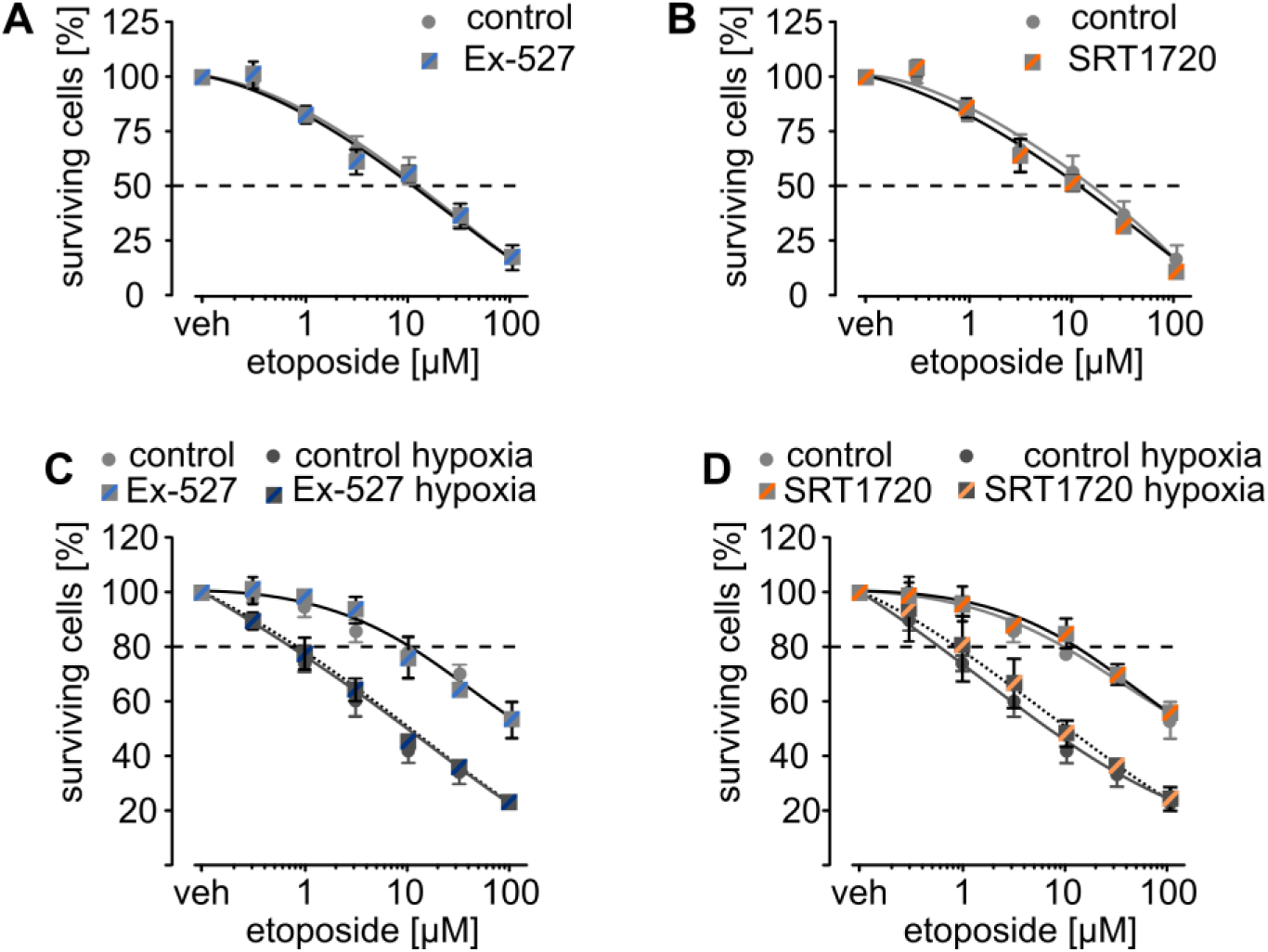
Modulation of SIRT1 activity has no impact on etoposide-induced cell death of NSPCs. NSPCs were treated with indicated concentrations of etoposide for 24 hours (**A,B**) or 1 h with 24 h recovery (**C,D**) with or without presence of Ex-527 (SIRT1 inhibition, **A,C**), SRT1720 (SIRT1 activation, **B,D**), or hypoxia (x % O_2_, **C,D**). Cell survival was determined via the CTB assay (n=3).

**Fig. S2.**
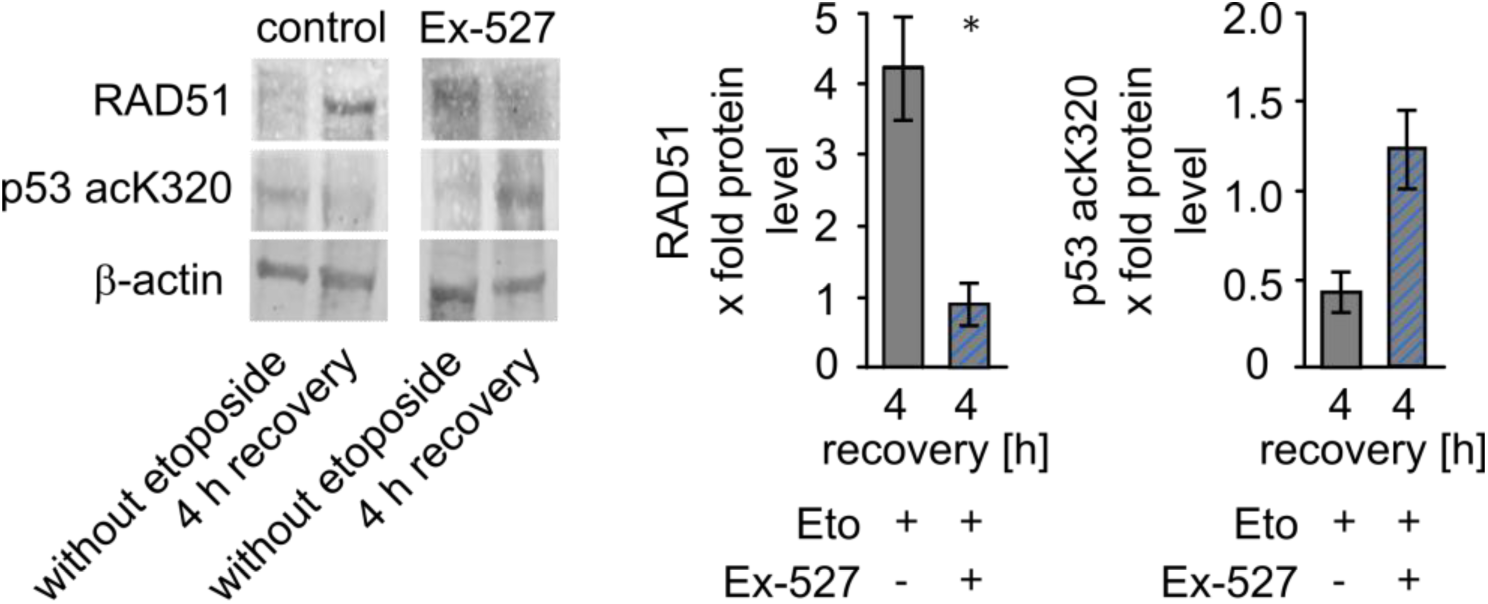
SIRT1 inhibition is linked to homologues recombination. NSPCs were treated with 1 µM od etoposide for 1 h ± EX-527. After 4 h of recovery, RAD51 (n=3) and acetylated p53K320 (n=2) were visualized by respective antibodies via western blots and the levels were quantified. Significance was determined by Student’s t-test, *: p<0.05.

## References

1. Alt, F. W., & Schwer, B. (2018). DNA double-strand breaks as drivers of neural genomic change, function, and disease. DNA Repair (Amst.), 71, 158–163.

2. Alt, F. W., Wei, P.-C., & Schwer, B. (2017). Recurrently breaking genes in neural progenitors: Potential roles of DNA breaks in neuronal function, degeneration and cancer. In Genome Editing in Neurosciences (pp. 63–72). Springer.

3. Arias-Lopez, C., Lazaro-Trueba, I., Kerr, P., Lord, C. J., Dexter, T., Iravani, M., Ashworth, A., & Silva, A. (2006). P53 modulates homologous recombination by transcriptional regulation of the RAD51 gene. EMBO Rep., 7(2), 219–224.

4. Barazzuol, L., Ju, L., & Jeggo, P. A. (2017). A coordinated DNA damage response promotes adult quiescent neural stem cell activation. PLoS Biology, 15(5), e2001264. 10.1371/journal.pbio.2001264

5. Bertrand, P., Saintigny, Y., & Lopez, B. S. (2004). P53’s double life: Transactivation-independent repression of homologous recombination. Trends Genet., 20(6), 235–243.

6. Bräutigam, L., Jensen, L. D. E., Poschmann, G., Nyström, S., Bannenberg, S., Dreij, K., Lepka, K., Prozorovski, T., Montano, S. J., Aktas, O., Uhlén, P., Stühler, K., Cao, Y., Holmgren, A., & Berndt, C. (2013). Glutaredoxin regulates vascular development by reversible glutathionylation of sirtuin 1. Proc. Natl. Acad. Sci. U. S. A., 110(50), 20057–20062.

7. Caito, S., Rajendrasozhan, S., Cook, S., Chung, S., Yao, H., Friedman, A. E., Brookes, P. S., & Rahman, I. (2010). SIRT1 is a redox-sensitive deacetylase that is post-translationally modified by oxidants and carbonyl stress. FASEB J., 24(9), 3145–3159.

8. Chang, H.-C., & Guarente, L. (2014). SIRT1 and other sirtuins in metabolism. Trends Endocrinol. Metab., 25(3), 138–145.

9. Chau, J. F. L., Jia, D., Wang, Z., Liu, Z., Hu, Y., Zhang, X., Jia, H., Lai, K. P., Leong, W. F., Au, B. J., Mishina, Y., Chen, Y.-G., Biondi, C., Robertson, E., Xie, D., Liu, H., He, L., Wang, X., Yu, Q., & Li, B. (2012). A crucial role for bone morphogenetic protein-Smad1 signalling in the DNA damage response. Nature Communications, 3, 836. 10.1038/ncomms1832

10. Cheng, H.-L., Mostoslavsky, R., Saito, S., Manis, J. P., Gu, Y., Patel, P., Bronson, R., Appella, E., Alt, F. W., & Chua, K. F. (2003). Developmental defects and p53 hyperacetylation in Sir2 homolog (SIRT1)-deficient mice. Proceedings of the National Academy of Sciences, 100(19), 10794– 10799. 10.1073/pnas.1934713100

11. De Filippis, L., & Delia, D. (2011). Hypoxia in the regulation of neural stem cells. Cell. Mol. Life Sci., 68(17), 2831–2844.

12. Delint-Ramirez, I., & Madabhushi, R. (2025). DNA damage and its links to neuronal aging and degeneration. Neuron, 113(1), 7–28. 10.1016/j.neuron.2024.12.001

13. Dobbin, M. M., Madabhushi, R., Pan, L., Chen, Y., Kim, D., Gao, J., Ahanonu, B., Pao, P.-C., Qiu, Y., Zhao, Y., & Tsai, L.-H. (2013). SIRT1 collaborates with ATM and HDAC1 to maintain genomic stability in neurons. Nat. Neurosci., 16(8), 1008–1015.

14. Fang, E. F., Kassahun, H., Croteau, D. L., Scheibye-Knudsen, M., Marosi, K., Lu, H., Shamanna, R. A., Kalyanasundaram, S., Bollineni, R. C., Wilson, M. A., Iser, W. B., Wollman, B. N., Morevati, M., Li, J., Kerr, J. S., Lu, Q., Waltz, T. B., Tian, J., Sinclair, D. A., … Bohr, V. A. (2016). NAD+ Replenishment Improves Lifespan and Healthspan in Ataxia Telangiectasia Models via Mitophagy and DNA Repair. Cell Metabolism, 24(4), 566–581. 10.1016/j.cmet.2016.09.004

15. Filipponi, D., Emelyanov, A., Muller, J., Molina, C., Nichols, J., & Bulavin, D. V. (2019). DNA damage signaling-induced cancer cell reprogramming as a driver of tumor relapse. Mol. Cell, 74(4), 651–663.e8.

16. Gleeson, J. G., Allen, K. M., Fox, J. W., Lamperti, E. D., Berkovic, S., Scheffer, I., Cooper, E. C., Dobyns, W. B., Minnerath, S. R., Ross, M. E., & Walsh, C. A. (1998). Doublecortin, a Brain-Specific Gene Mutated in Human X-Linked Lissencephaly and Double Cortex Syndrome, Encodes a Putative Signaling Protein. Cell, 92(1), 63–72. 10.1016/S0092-8674(00)80899-5

17. Guo, W., Patzlaff, N. E., Jobe, E. M., & Zhao, X. (2012). Isolation of multipotent neural stem or progenitor cells from both the dentate gyrus and subventricular zone of a single adult mouse. Nature Protocols, 7(11), 2005–2012. 10.1038/nprot.2012.123

18. Guo, W., Qian, L., Zhang, J., Zhang, W., Morrison, A., Hayes, P., Wilson, S., Chen, T., & Zhao, J. (2011). Sirt1 overexpression in neurons promotes neurite outgrowth and cell survival through inhibition of the mTOR signaling. J. Neurosci. Res., 89(11), 1723–1736.

19. Habich, M., & Riemer, J. (2017). Detection of Cysteine Redox States in Mitochondrial Proteins in Intact Mammalian Cells. In D. Mokranjac & F. Perocchi (Eds.), Mitochondria (Vol. 1567, pp. 105–138). Springer New York. 10.1007/978-1-4939-6824-4_8

20. Hajji, N., Wallenborg, K., Vlachos, P., Füllgrabe, J., Hermanson, O., & Joseph, B. (2010). Opposing effects of hMOF and SIRT1 on H4K16 acetylation and the sensitivity to the topoisomerase II inhibitor etoposide. Oncogene, 29(15), 2192–2204.

21. Herranz, D., Muñoz-Martin, M., Cañamero, M., Mulero, F., Martinez-Pastor, B., Fernandez-Capetillo, O., & Serrano, M. (2010). Sirt1 improves healthy ageing and protects from metabolic syndrome-associated cancer. Nature Communications, 1, 3. 10.1038/ncomms1001

22. Homma, H., Tanaka, H., Jin, M., Jin, X., Huang, Y., Yoshioka, Y., Bertens, C. J., Tsumaki, K., Kondo, K., Shiwaku, H., Tagawa, K., Akatsu, H., Atsuta, N., Katsuno, M., Furukawa, K., Ishiki, A., Waragai, M., Ohtomo, G., Iwata, A., … Okazawa, H. (2021). DNA damage in embryonic neural stem cell determines FTLDs’ fate via early-stage neuronal necrosis. Life Sci. Alliance, 4(7), e202101022.

23. Jackson, S. P., & Bartek, J. (2009). The DNA-damage response in human biology and disease. Nature, 461(7267), 1071–1078. 10.1038/nature08467

24. Knights, C. D., Catania, J., Di Giovanni, S., Muratoglu, S., Perez, R., Swartzbeck, A., Quong, A. A., Zhang, X., Beerman, T., Pestell, R. G., & Avantaggiati, M. L. (2006). Distinct p53 acetylation cassettes differentially influence gene-expression patterns and cell fate. The Journal of Cell Biology, 173(4), 533–544. 10.1083/jcb.200512059

25. Konopka, A., Atkin, J. D., & Mitra, J. (2022). Editorial: Defective DNA damage response-Repair axis in post-mitotic neurons in human health and neurodegenerative diseases. Frontiers in Cellular Neuroscience, 16, 1009760. 10.3389/fncel.2022.1009760

26. Krishna, S., Wagener, B. M., Liu, H. P., Lo, Y.-C., Sterk, R., Petrini, J. H. J., & Nickoloff, J. A. (2007). Mre11 and Ku regulation of double-strand break repair by gene conversion and break-induced replication. DNA Repair (Amst.), 6(6), 797–808.

27. Kweon, K. H., Lee, C. R., Jung, S. J., Ban, E. J., Kang, S.-W., Jeong, J. J., Nam, K.-H., Jo, Y. S., Lee, J., & Chung, W. Y. (2014). Sirt1 induction confers resistance to etoposide-induced genotoxic apoptosis in thyroid cancers. Int. J. Oncol., 45(5), 2065–2075.

28. Kwon, J., Lee, S., Kim, Y.-N., & Lee, I. H. (2019). Deacetylation of CHK2 by SIRT1 protects cells from oxidative stress-dependent DNA damage response. Exp. Mol. Med., 51(3), 1–9.

29. Lagunas-Rangel, F. A. (2019). Current role of mammalian sirtuins in DNA repair. DNA Repair (Amst.), 80, 85–92.

30. Lee, J.-S., Park, J.-R., Kwon, O.-S., Lee, T.-H., Nakano, I., Miyoshi, H., Chun, K.-H., Park, M.-J., Lee, H. J., Kim, S. U., & Cha, H.-J. (2015). SIRT1 is required for oncogenic transformation of neural stem cells and for the survival of “cancer cells with neural stemness” in a p53-dependent manner. Neuro. Oncol., 17(1), 95–106.

31. Lee, Y., Barnes, D. E., Lindahl, T., & McKinnon, P. J. (2000). Defective neurogenesis resulting from DNA ligase IV deficiency requires Atm. Genes Dev., 14(20), 2576–2580.

32. Lei, T., Du, S., Peng, Z., & Chen, L. (2022). Multifaceted regulation and functions of 53BP1 in NHEJ-mediated DSB repair (Review). International Journal of Molecular Medicine, 50(1), 90. 10.3892/ijmm.2022.5145

33. Lim, D. A., & Alvarez-Buylla, A. (2016). The adult ventricular-subventricular zone (V-SVZ) and olfactory bulb (OB) neurogenesis. Cold Spring Harb. Perspect. Biol., 8(5).

34. Liu, D. J., Hammer, D., Komlos, D., Chen, K. Y., Firestein, B. L., & Liu, A. Y.-C. (2014). SIRT1 knockdown promotes neural differentiation and attenuates the heat shock response. J. Cell. Physiol., 229(9), 1224–1235.

35. Mah, L.-J., El-Osta, A., & Karagiannis, T. C. (2010). GammaH2AX: a sensitive molecular marker of DNA damage and repair. Leukemia, 24(4), 679–686.

36. Milne, J. C., Lambert, P. D., Schenk, S., Carney, D. P., Smith, J. J., Gagne, D. J., Jin, L., Boss, O., Perni, R. B., Vu, C. B., Bemis, J. E., Xie, R., Disch, J. S., Ng, P. Y., Nunes, J. J., Lynch, A. V., Yang, H., Galonek, H., Israelian, K., … Westphal, C. H. (2007). Small molecule activators of SIRT1 as therapeutics for the treatment of type 2 diabetes. Nature, 450(7170), 712–716.

37. Mishra, P., Mittal, A. K., Kalonia, H., Madan, S., Ghosh, S., Sinha, J. K., & Rajput, S. K. (2021). SIRT1 Promotes Neuronal Fortification in Neurodegenerative Diseases through Attenuation of Pathological Hallmarks and Enhancement of Cellular Lifespan. Current Neuropharmacology, 19(7), 1019–1037. 10.2174/1570159X18666200729111744

38. Moretton, A., & Loizou, J. I. (2020). Interplay between cellular metabolism and the DNA damage response in cancer. Cancers (Basel), 12(8), 2051.

39. Murillo, L. C., Sutachan, J. J., & Albarracín, S. L. (2023). An update on neurobiological mechanisms involved in the development of chemotherapy-induced cognitive impairment (CICI). Toxicology Reports, 10, 544–553. 10.1016/j.toxrep.2023.04.015

40. Napper, A. D., Hixon, J., McDonagh, T., Keavey, K., Pons, J.-F., Barker, J., Yau, W. T., Amouzegh, P., Flegg, A., Hamelin, E., Thomas, R. J., Kates, M., Jones, S., Navia, M. A., Saunders, J. O., DiStefano, P. S., & Curtis, R. (2005). Discovery of indoles as potent and selective inhibitors of the deacetylase SIRT1. J. Med. Chem., 48(25), 8045–8054.

41. Ng, N., Purshouse, K., Foskolou, I. P., Olcina, M. M., & Hammond, E. M. (2018). Challenges to DNA replication in hypoxic conditions. FEBS J., 285(9), 1563–1571.

42. Oberdoerffer, P., Michan, S., McVay, M., Mostoslavsky, R., Vann, J., Park, S.-K., Hartlerode, A., Stegmuller, J., Hafner, A., Loerch, P., Wright, S. M., Mills, K. D., Bonni, A., Yankner, B. A., Scully, R., Prolla, T. A., Alt, F. W., & Sinclair, D. A. (2008). SIRT1 redistribution on chromatin promotes genomic stability but alters gene expression during aging. Cell, 135(5), 907–918.

43. Orhan, E., Velazquez, C., Tabet, I., Sardet, C., & Theillet, C. (2021). Regulation of RAD51 at the transcriptional and functional levels: What prospects for cancer therapy? Cancers (Basel), 13(12), 2930.

44. Prozorovski, T., Ingwersen, J., Lukas, D., Göttle, P., Koop, B., Graf, J., Schneider, R., Franke, K., Schumacher, S., Britsch, S., Hartung, H.-P., Küry, P., Berndt, C., & Aktas, O. (2019). Regulation of sirtuin expression in autoimmune neuroinflammation: Induction of SIRT1 in oligodendrocyte progenitor cells. Neurosci. Lett., 704, 116–125.

45. Prozorovski, T., Schneider, R., Berndt, C., Hartung, H.-P., & Aktas, O. (2015). Redox-regulated fate of neural stem progenitor cells. Biochimica Et Biophysica Acta, 1850(8), 1543–1554. 10.1016/j.bbagen.2015.01.022

46. Prozorovski, T., Schulze-Topphoff, U., Glumm, R., Baumgart, J., Schröter, F., Ninnemann, O., Siegert, E., Bendix, I., Brüstle, O., Nitsch, R., Zipp, F., & Aktas, O. (2008). Sirt1 contributes critically to the redox-dependent fate of neural progenitors. Nat. Cell Biol., 10(4), 385–394.

47. Qing, X., Zhang, G., & Wang, Z. (2023). DNA damage response in neurodevelopment and neuromaintenance. The FEBS Journal, 290(13), 3300–3310. 10.1111/febs.16535

48. Rafalski, V. A., Ho, P. P., Brett, J. O., Ucar, D., Dugas, J. C., Pollina, E. A., Chow, L. M. L., Ibrahim, A., Baker, S. J., Barres, B. A., Steinman, L., & Brunet, A. (2013). Expansion of oligodendrocyte progenitor cells following SIRT1 inactivation in the adult brain. Nat. Cell Biol., 15(6), 614–624.

49. Raineteau, O., Rietschin, L., Gradwohl, G., Guillemot, F., & Gähwiler, B. H. (2004). Neurogenesis in hippocampal slice cultures. Mol. Cell. Neurosci., 26(2), 241–250.

50. Rasti, G., Becker, M., Vazquez, B. N., Espinosa-Alcantud, M., Fernández-Duran, I., Gámez-García, A., Ianni, A., Gonzalez, J., Bosch-Presegué, L., Marazuela-Duque, A., Guitart-Solanes, A., Segura-Bayona, S., Bech-Serra, J.-J., Scher, M., Serrano, L., Shankavaram, U., Erdjument-Bromage, H., Tempst, P., Reinberg, D., … Vaquero, A. (2023). SIRT1 regulates DNA damage signaling through the PP4 phosphatase complex. Nucleic Acids Research, 51(13), 6754–6769. 10.1093/nar/gkad504

51. Ren, J., Wang, X., Dong, C., Wang, G., Zhang, W., Cai, C., Qian, M., Yang, D., Ling, B., Ning, K., Mao, Z., Liu, B., Wang, T., Xiong, L., Wang, W., Liang, A., Gao, Z., & Xu, J. (2022). Sirt1 Protects Subventricular Zone-Derived Neural Stem Cells from DNA Double-Strand Breaks and Contributes to Olfactory Function Maintenance in Aging Mice. Stem Cells, 40(5), 493–507. 10.1093/stmcls/sxac008

52. Ribeiro, J. H., Altinisik, N., Rajan, N., Verslegers, M., Baatout, S., Gopalakrishnan, J., & Quintens, R. (2023). DNA damage and repair: Underlying mechanisms leading to microcephaly. Frontiers in Cell and Developmental Biology, 11, 1268565. 10.3389/fcell.2023.1268565

53. Rimmelé, P., Bigarella, C. L., Liang, R., Izac, B., Dieguez-Gonzalez, R., Barbet, G., Donovan, M., Brugnara, C., Blander, J. M., Sinclair, D. A., & Ghaffari, S. (2014). Aging-like phenotype and defective lineage specification in SIRT1-deleted hematopoietic stem and progenitor cells. Stem Cell Reports, 3(1), 44–59.

54. Saharan, S., Jhaveri, D. J., & Bartlett, P. F. (2013). SIRT1 regulates the neurogenic potential of neural precursors in the adult subventricular zone and hippocampus. J. Neurosci. Res., 91(5), 642–659.

55. Satoh, A., Brace, C. S., Rensing, N., Cliften, P., Wozniak, D. F., Herzog, E. D., Yamada, K. A., & Imai, S.-I. (2013). Sirt1 extends life span and delays aging in mice through the regulation of Nk2 homeobox 1 in the DMH and LH. Cell Metabolism, 18(3), 416–430. 10.1016/j.cmet.2013.07.013

56. Schneider, R., Koop, B., Schröter, F., Cline, J., Ingwersen, J., Berndt, C., Hartung, H.-P., Aktas, O., & Prozorovski, T. (2016). Activation of Wnt signaling promotes hippocampal neurogenesis in experimental autoimmune encephalomyelitis. Mol. Neurodegener., 11(1), 53.

57. Shimada, M., Dumitrache, L. C., Russell, H. R., & McKinnon, P. J. (2015). Polynucleotide kinase-phosphatase enables neurogenesis via multiple DNA repair pathways to maintain genome stability. EMBO J., 34(19), 2465–2480.

58. Shull, E. R. P., Lee, Y., Nakane, H., Stracker, T. H., Zhao, J., Russell, H. R., Petrini, J. H. J., & McKinnon, P. J. (2009). Differential DNA damage signaling accounts for distinct neural apoptotic responses in ATLD and NBS. Genes Dev., 23(2), 171–180.

59. Solomon, J. M., Pasupuleti, R., Xu, L., McDonagh, T., Curtis, R., DiStefano, P. S., & Huber, L. J. (2006). Inhibition of SIRT1 catalytic activity increases p53 acetylation but does not alter cell survival following DNA damage. Mol. Cell. Biol., 26(1), 28–38.

60. Song, J., Christian, K. M., Ming, G.-L., & Song, H. (2012). Modification of hippocampal circuitry by adult neurogenesis. Dev. Neurobiol., 72(7), 1032–1043.

61. Sun, Z., Zhao, S., Suo, X., & Dou, Y. (2022). Sirt1 protects against hippocampal atrophy and its induced cognitive impairment in middle-aged mice. BMC Neuroscience, 23(1), 33. 10.1186/s12868-022-00718-8

62. Thapa, R., Moglad, E., Afzal, M., Gupta, G., Bhat, A. A., Hassan Almalki, W., Kazmi, I., Alzarea, S. I., Pant, K., Singh, T. G., Singh, S. K., & Ali, H. (2024). The role of sirtuin 1 in ageing and neurodegenerative disease: A molecular perspective. Ageing Research Reviews, 102, 102545. 10.1016/j.arr.2024.102545

63. Torre, M., Dey, A., Woods, J. K., & Feany, M. B. (2021). Elevated Oxidative Stress and DNA Damage in Cortical Neurons of Chemotherapy Patients. Journal of Neuropathology and Experimental Neurology, 80(7), 705–712. 10.1093/jnen/nlab074

64. Wagle, R., & Song, Y.-H. (2020). Ionizing radiation reduces larval brain size by inducing premature differentiation of Drosophila neural stem cells. Biochemical and Biophysical Research Communications, 523(3), 555–560. 10.1016/j.bbrc.2019.12.047

65. Wang, R.-H., Sengupta, K., Li, C., Kim, H.-S., Cao, L., Xiao, C., Kim, S., Xu, X., Zheng, Y., Chilton, B., Jia, R., Zheng, Z.-M., Appella, E., Wang, X. W., Ried, T., & Deng, C.-X. (2008). Impaired DNA damage response, genome instability, and tumorigenesis in SIRT1 mutant mice. Cancer Cell, 14(4), 312–323.

66. Wang, W., Esbensen, Y., Kunke, D., Suganthan, R., Rachek, L., Bjørås, M., & Eide, L. (2011). Mitochondrial DNA damage level determines neural stem cell differentiation fate. The Journal of Neuroscience: The Official Journal of the Society for Neuroscience, 31(26), 9746–9751. 10.1523/JNEUROSCI.0852-11.2011

67. Wei, P.-C., Chang, A. N., Kao, J., Du, Z., Meyers, R. M., Alt, F. W., & Schwer, B. (2016). Long neural genes harbor recurrent DNA break clusters in neural stem/progenitor cells. Cell, 164(4), 644– 655.

68. Xu, X., An, H., Wu, C., Sang, R., Wu, L., Lou, Y., Yang, X., & Xi, Y. (2023). HR repair pathway plays a crucial role in maintaining neural stem cell fate under irradiation stress. Life Science Alliance, 6(8), e202201802. 10.26508/lsa.202201802

69. Zhang, W., Wu, H., Yang, M., Ye, S., Li, L., Zhang, H., Hu, J., Wang, X., Xu, J., & Liang, A. (2016). SIRT1 inhibition impairs non-homologous end joining DNA damage repair by increasing Ku70 acetylation in chronic myeloid leukemia cells. Oncotarget, 7(12), 13538–13550.

